# Building a learnable universal coordinate system for single-cell atlas with a joint-VAE model

**DOI:** 10.1101/2021.09.09.459281

**Authors:** Haoxiang Gao, Kui Hua, Lei Wei, Xinze Wu, Sijie Chen, Qijin Yin, Rui Jiang, Xuegong Zhang

## Abstract

A universal coordinate system that can ensemble the huge number of cells and capture their heterogeneities is of vital importance for constructing large-scale cell atlases as references for future molecular and cellular studies. Studies have shown that cells in complex organs exhibit multifaceted heterogeneities in their transcriptomic features at multiple resolutions. This nature of complexity makes it hard to design a fixed coordinate system through a combination of known features. It is desirable to build a learnable universal coordinate model that can capture major heterogeneities and serve as a controlled generative model for data argumentation. We developed UniCoord, a specially tuned joint-VAE model to represent single-cell transcriptomic data in a lower-dimensional latent space with high interpretability. Each latent dimension can represent either discrete or continuous feature, and either supervised by prior knowledge or unsupervised. The original transcriptomic profiles can be regenerated from the latent dimensions. The latent dimensions can be easily reconfigured to generate transcriptomic profiles of pseudo cells with desired properties. UniCoord can also be used as a pre-trained model to analyze new data with unseen cell types and thus can serve as a feasible framework for cell annotation and comparison. UniCoord provides a prototype for a learnable universal coordinate framework to enable better analysis and generation of cells with highly orchestrated functions and heterogeneities.

## Introduction

Cells in complex organs exhibit multifaceted heterogeneities, determining the various physiological and pathological phenomena of life. Since the discovery of cells, researchers have always been trying to classify cells into different cell types with their morphological features, molecular markers or cellular functions^1^. With the rapid development of single-cell omics technology, there arises the ambition to build cell atlases that can serve as a reference to describe the multifaceted heterogeneities of cells^2–4^. When given a cell, a desired cell atlas should be able to locate the cell to a specific body position and differentiation stage by assigning spatial and temporal coordinates. Moreover, the atlas should describe cell types/states as well as the activities of various biological processes of the cell, all of which can be summarized as functional coordinates. A universal coordinate system is essential to achieve this goal. A well-designed universal coordinate system can organize the huge number of cells within a cell atlas in a quantitative way, and thus benefit future molecular and cellular studies.

Many studies have been proposed to quantify the spatial, temporal, and functional features of cells. For example, a variation of tools have been developed to construct temporal trajectories or assign a pseudo-time score to each cell in single-cell RNA-seq (scRNA-seq) data^5–10^. Similarly, with the rapid development of spatial profiling technologies, tools keep emerging to infer cell positions for scRNA-seq data^11–15^. Diverse features or systems were proposed to illustrate the multifaceted functional characteristics of cells, such as hierarchically organized cell types^4,16,17^, the continuum of cell states related to tumor progression^18,19^ and cell cycle^20^, and the index of macrophage activation states^21^. Beyond these approaches focusing on specialized cellular features, some works attempted to build information systems that organize these features within a unified framework, such as a spatial coordinate system that labels the original sampling site of cells^22^, and methods that embed transcriptomic profiles of cells into a latent space without explicit interpretations^23,24^.

All these attempts tried to describe cells with specific features, which can hardly serve as a universal coordinate system as they don’t match the complex nature of cells. For example, anatomic structures of human bodies are conserved in the population, but there are great variations and flexibilities in morphology and sizes among individuals, letting alone the cellular conformation^25^. Studies have shown that cells exhibit multifaceted heterogeneities in their transcriptomic features, including spatial, temporal and functional gradients at multiple resolutions^26–30^. This makes it hard to design a fixed coordinate system with known cell types, locations and sampling time points to capture and index all the gradients, especially when considering the fact that the currently measured features are still far from providing the whole information of cells. It is desirable to build a learnable universal coordinate system that can capture all major heterogeneities in currently available data and can be compatible for future extensions when data with richer information are available. Such a system should be able to integrate both discrete and continuous coordinates within a single model, and these coordinates are preferably with interpretability. The system should also provide possibilities of generating pseudo cells by reconfiguring coordinates to help explore cell states that are not included in existing data or can hardly be observed by experimental approaches.

In this work, we developed a Universal Coordinate model (UniCoord) that learns to represent cells with a series of discrete and continuous features according to transcriptomic profiles. It used a specially-tuned joint variational autoencoder (VAE) model to learn key features that best represent cellular heterogeneities. Each feature can be either discrete or continuous, and either supervised by prior knowledge or unsupervised. We applied UniCoord on several datasets, and the results showed that UniCoord is able to capture multiple key cellular features such as spatial, temporal and functional gradients from massive data. These features are powerful for accurate data reconstruction and label identification. Furthermore, UniCoord can serve as a controlled generative model for data argumentation, such as generating pseudo cells with desired features and interpolating extra cells in spatial or temporal gradients to fill the gaps between sampled cell states. UniCoord can be used as a pre-trained model feasible to analyze new data with unseen cell types, such as using a UniCoord model trained by healthy data to analyze disease data. UniCoord provides a prototype for a learnable universal coordinate framework for analyzing the highly orchestrated functions and multifaceted heterogeneities of diverse cells, and paves the way towards a seamless cell atlas by unified organization and data argumentation.

## Results

### Overview of UniCoord

We developed UniCoord to learn key features that represent cellular heterogeneities from scRNA-seq data. We devised a specially tuned joint-VAE model to represent the transcriptomic profile of a single cell in a low-dimensional latent space. A conventional VAE model^31^ includes an encoder and a decoder (Fig. 1a, see Methods for details). The encoder transforms the transcriptomic profile of a single cell into a set of means to represent the features about cellular heterogeneities, and a set of variances to deal with uncertainty. The decoder samples from the latent space according to the distribution defined by the means and variances learned by the encoder, and then transformed the sample into a generated transcriptomic profile.

**Figure 1.**
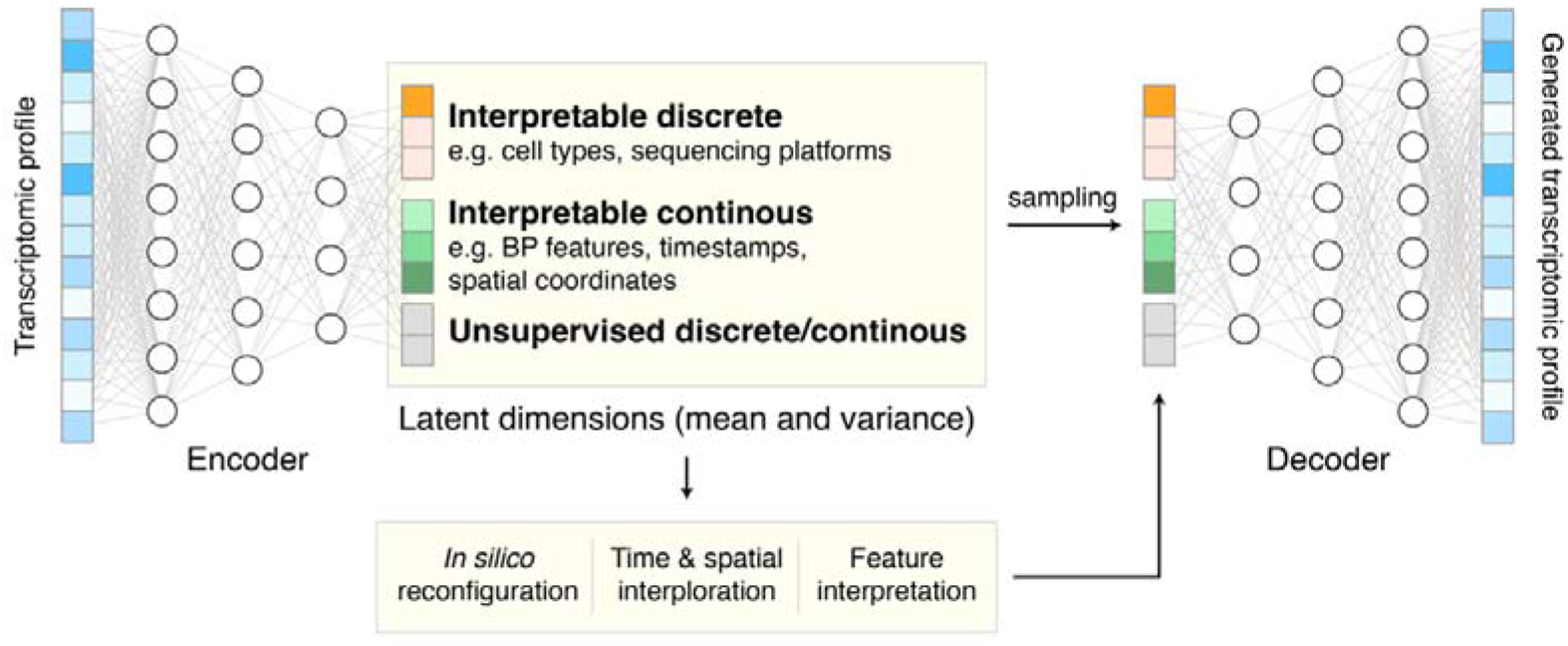
The schematic diagram of UniCoord.

The goal of UniCoord is to represent the transcriptomic profiles of cells in the latent space with interpretability. We thus designed each dimension in the latent space to be either supervised by prior knowledge or unsupervised. We trained the supervised dimensions to capture information corresponding to well defined features of the cell, such as the activity of a biological pathway, the differentiation stage, or the clinical diagnoses of the cell’s donor. We trained the unsupervised dimensions to capture complementary, yet unknown information of the cell, such as an unsupervised classification or score of cells. Through this approach, the latent dimensions can be regarded as the universal coordinates of cells.

Both discrete and continuous features can be used to represent cellular heterogeneities. For instance, the cell type and sequencing platform are commonly-considered discrete features in RNA-seq studies. The spatial, temporal and functional gradients are continuous features of vital importance to the analysis of biological processes. We proposed a model based on joint-VAE^32^, a disentangled representation framework that can deal with both discrete and continuous features in a single model, to handle these multifaceted heterogeneities. We considered two main aspects of loss in model training. We considered the reconstruction loss between the original and constructed data, which guaranteed the accuracy of the VAE model. Besides, we designed different loss functions for different forms of supervised features to make these features consistent with prior knowledge (see Methods for details).

### Generating pseudo single-cell data by *in silico* reconfiguration

Abundant transcriptomic profiles of cells are essential in the analysis of physiological and pathological processes. However, the cell numbers in existing data are often not sufficient, and it is challenging to experimentally observe certain cell types or states, especially for intermediate states. With UniCoord, we can reconfigure any latent dimension into a desired value and then obtain pseudo single-cell transcriptomic profile data. We named this procedure as *in silico* reconfiguration.

We first evaluated *in silico* reconfiguration of discrete features, such as the sequencing platforms and cell types. We trained a UniCoord model with the lung data in hECA, and used cell types, sequencing platforms, and unsupervised continuous latent dimensions as the latent dimensions. The original scRNA-seq data derived from different sequencing platforms were separated in the Uniform Manifold Approximation and Projection (UMAP) plot (Fig. 2a), which is mainly due to the distinct distribution of data such as the median number of expressed genes in each cell (Fig. S1a). We used the latent representation of original cells as the seed and reconfigured the mean value of the latent dimension corresponding to the sequencing platform into “10X”. We then sampled random variables from the modified distribution, and used the decoder to generate pseudo transcriptomic profiles according to these sampled random variables. We found that the generated cells were joined together and clustered well by cell types (Fig. 2b). These generated cells showed similar distributions of the number of expressed genes, regardless the platform that their corresponding original cells were derived from (Fig. S1b). We then reconfigured the cell types of all cells into a specified one. When we reconfigured the cell type as T cells and the sequencing platform as “10X”, the generated data highly expressed markers of T cells, including PTPRC, CD3E, and TRAC (Fig. 2c-d). When we reconfigured the cell type as fibrocytes and the sequencing platform as “10X”, these immune-related markers were turned off, and extracellular matrix markers such as DCN and COL1A1 were highly expressed (Fig. 2e-f).

**Figure 2.**
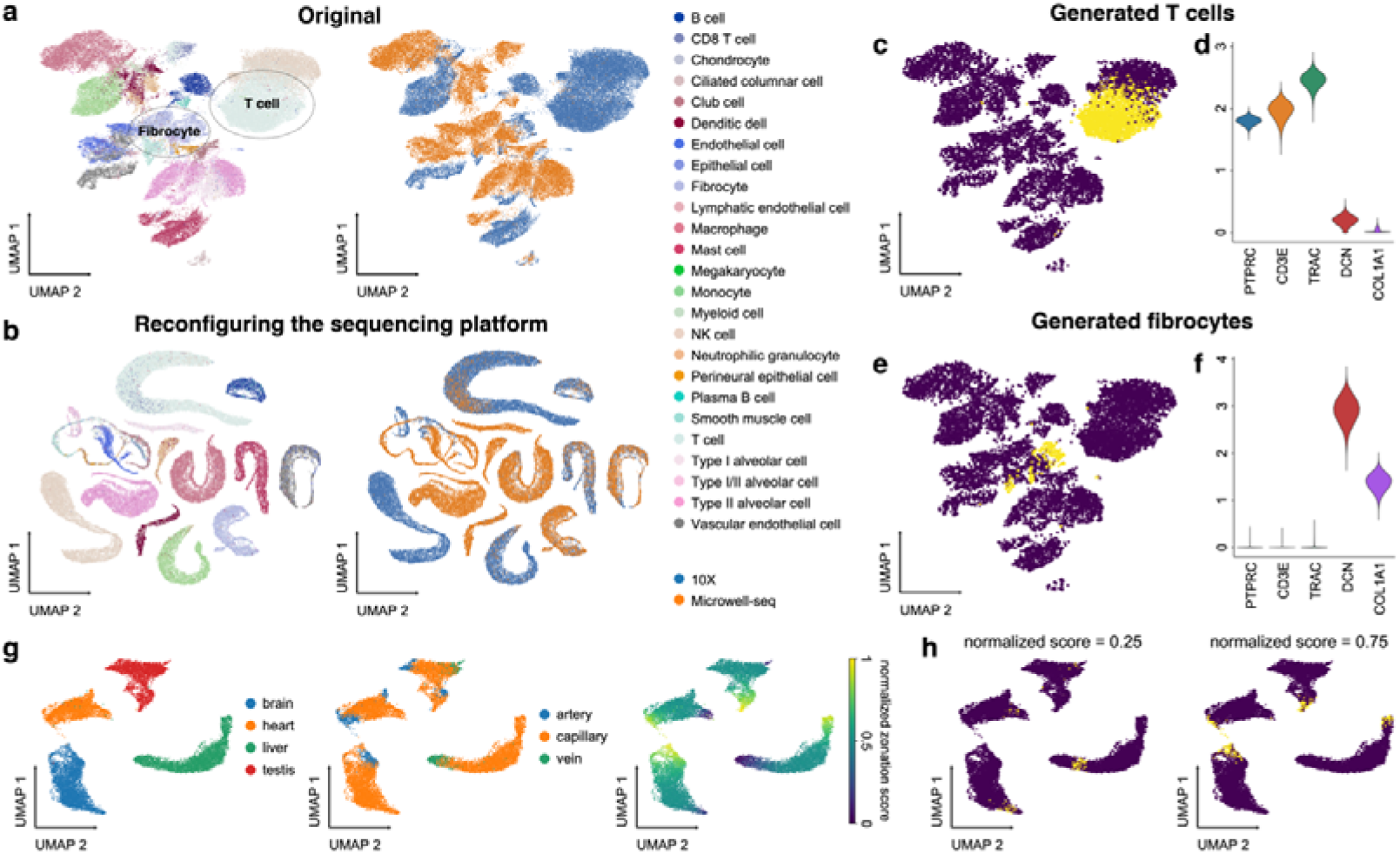
*In silico* reconfiguration with UniCoord generates cells with designated features. (a) UMAP plots showing the original hECA lung cells colored by cell types (left) or sequencing platforms (right). (b) UMAP plots showing the generated hECA lung cells with the sequencing platform reconfigured into “10X”, colored by cell types (left) and original sequencing platforms (right). (c) The UMAP plot of original cells (blue) and cells with the cell type reconfigured into T cells (yellow). (d) Expression levels of T cell and fibrocyte markers in cells with the cell type reconfigured into T cells. (e) The UMAP plot of original cells (blue) and cells with the cell type reconfigured into fibrocytes (yellow). (f) Expression levels of T cell and fibrocyte markers in cells with the cell type reconfigured into fibrocytes. (g) Vascular endothelial cells from four mouse organs, colored by tissues (left), vessel types (middle), or normalized artery-capillary-vein zonation scores (right). The zonation scores were normalized to be between 0 and 1. (h) UMAP plots of original cells (blue) and cells with zonation scores reconfigured into different values (yellow). For both models, all genes in the dataset were adopted to perform the experiments, and the number of unsupervised latent dimensions was set as 50. For (a) and (b), all genes were used for visualization. For (c), (e), (g) and (h), highly variable genes (HVGs) were used for visualization. For (c), (e) and (h), for each generated cell, we calculated its nearest neighbor in the original dataset for visualization.

*In silico* reconfiguration can also be applied to continuous features. To demonstrate this, we trained a UniCoord model with the data of vascular endothelial cells from four tissues (brain, heart, liver, and testis) in a mouse endothelial atlas^33^. The original study discovered an artery-capillary-vein trajectory in vascular endothelial cells and raised a zonation score to describe the location of a cell on the trajectory. Thus, we used the information of tissues, zonation scores (normalized to be between 0 and 1), and unsupervised continuous latent dimensions as the latent dimensions. The result showed that UniCoord successfully reconstructed the information of tissues as well as the zonation trajectory (Fig. 2g). We then reconfigured the zonation scores of all cells and generated pseudo vascular endothelial cells. The generated cells were accurately placed at the designated locations on the trajectory (Fig. 2h and Movie S1). These results demonstrated the advantage of *in silico* reconfiguration with UniCoord in generating pseudo cells with desired properties, which could be helpful to analyze and integrate datasets from different sources.

### Interpolating timestamps or spatial coordinates to fill data gaps

It is impossible to obtain continuous observation of cellular transcriptomic profiles through high-throughput sequencing technologies. One common approach to obtain time-series single-cell transcriptomic data is to measure samples at different time points. However, there still exist inevitable gaps between time points. We used the *in-silico* reconfiguration approach to fill these gaps by timestamp interpolation. We trained a UniCoord model with a mouse embryo dataset^34^. In this dataset, cells were treated with doxycycline to induce mouse embryo fibroblasts (MEFs) de-differentiating into iPSCs, and were then transferred to either serum-free 2i medium or serum medium on day 8 (Fig. 3a). Cells in the serum medium are more likely to re-differentiate into stromal cells and neurons (Fig. 3b-c). The dataset covered a total of 18 days at half-day intervals, forming a discrete trajectory (Fig. 3a). We used timestamps and unsupervised continuous latent dimensions as the latent dimensions. After training, we sampled cells from each time point and used their latent representations as seeds. We reconfigured the latent dimension denoting the timestamp of each sampled cell by adding the original value with a uniformly random variable between –0.5 to 0.5 day. Through this approach, we interpolated the missing time points and created a continuous trajectory (Fig. 3d). We found that the trend in the evolution of cell types became more evident (Fig. 3f). Furthermore, though we didn’t encode any information about the experimental design, the interpolated results showed clear subgroups of cells with different treatments (Fig. 3e). The results demonstrated that UniCoord is feasible to discover and preserve the information about cell state evolution and fate decision.

**Figure 3.**
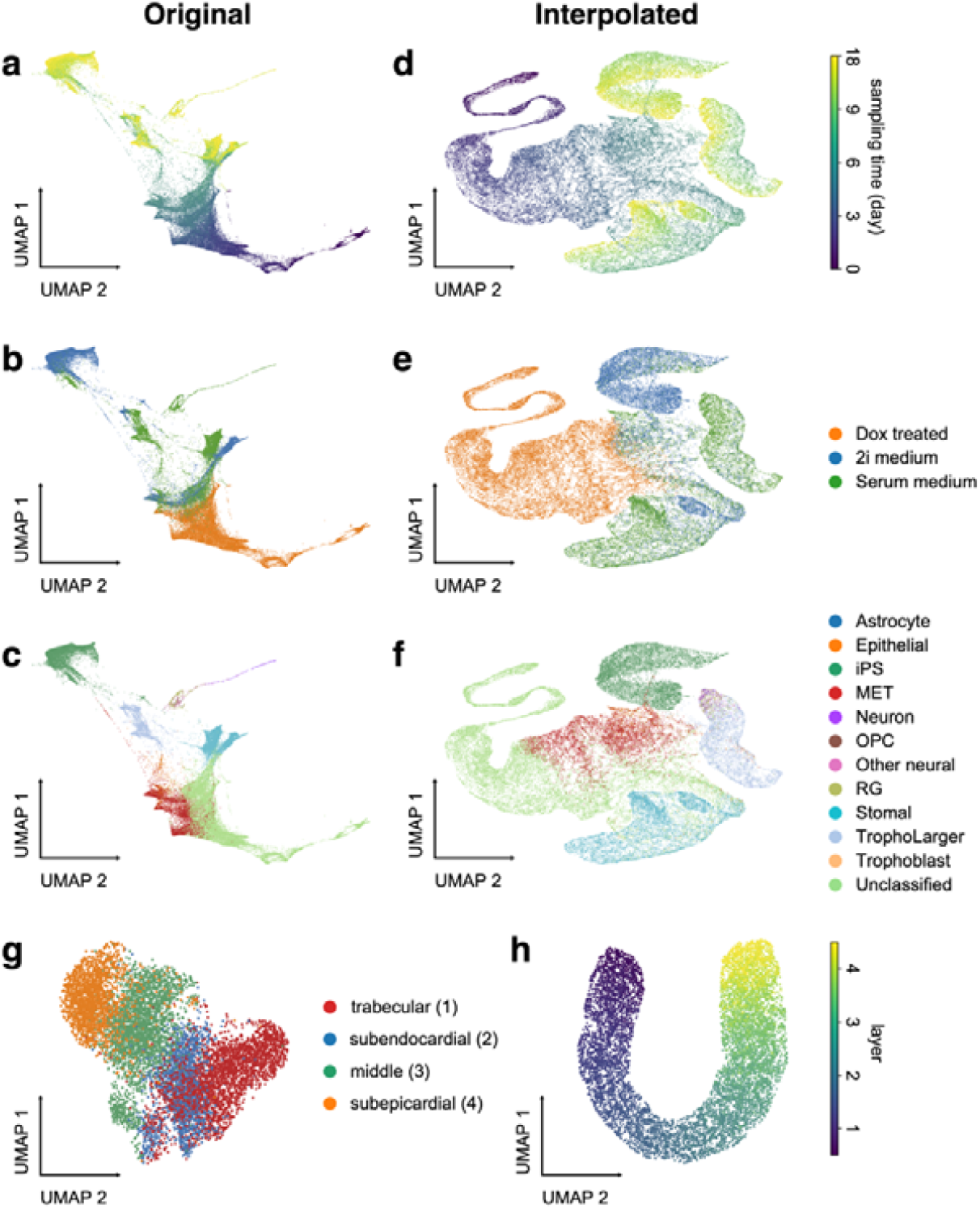
UniCoord interpolated discrete timestamps or spatial coordinates into continuous trajectories. (a-c) Mouse embryo data visualized by the low-dimensional visualization provided by the original study, cells colored by sampling days (a), treatments (b), and cell types (c). (d-f) UniCoord-interpolated mouse embryo data, cells colored by continuous sampling time (d), treatments (e), or cell types (f). Dox: doxycycline; iPS: induced pluripotent stem cells; MET: cells undergoing a mesenchymal-to-epithelial transition; OPC: oligodendrocyte precursor cells; RG: radial glial cells. (g-h) UMAP plots showing real (g) and interpolated (h) CMs from the left ventricle. The corresponding layer number was shown in the legend of (g). For both models, all genes in the dataset were adopted to perform the experiments, and the number of unsupervised latent dimensions was set as 50. HVGs were used for visualization all UMAP plots.

UniCoord can also be applied to reconstruct spatial trajectories by interpolating spatial coordinates. In our recent work on human heart cell atlas^28^, we sampled cardiomyocytes (CMs) from four layers of the left ventricle with different depths, and found these CMs from different layers exhibited distinct characteristics (Fig. 3g). We used UniCoord to interpolate the continuous change between these layers. We trained a UniCoord model with the data using the information of sample layers and unsupervised continuous latent dimensions as the latent dimensions. After training, we sampled cells from each layer, and reconfigured the latent dimension denoting the layer information by adding the original value with a uniformly random variable between –1 to 1. We generated pseudo cells with this approach and found that the generated data formed a continuous spatial trajectory (Fig. 3h). All the results demonstrated the versatility and potential of UniCoord in reconstructing spatial and temporal trajectories by interpolating timestamps or spatial coordinates, allowing for more comprehensive and accurate analyses of complex biological processes.

### Pre-training UniCoord model with cell atlas data for analyzing disease-related cells

We applied UniCoord on the human Ensembled Cell Atlas (hECA) data^4^ which has a total of 1.09 million cells to represent the cellular heterogeneity in the atlas. We randomly sampled 50,000 cells from the dataset and used these cells to train a UniCoord model. We used three aspects as the latent dimensions: cell types, sequencing platforms, and biological process (BP) features that represent the information of specific biological processes. To calculate BP features, we first applied AUCell^35^ on the transcriptomic profiles of the training cells to convert gene expression levels into the activity strengths of Gene Ontology Biological Process (GOBP) terms^36,37^. We then trained a random forest model that used these strengths to classify cell types, and identified the top 100 GOBP terms with the highest important scores. After, we clustered these terms into 30 groups and selected the term that showed the activity strength in each group. These selected GOBP terms were regarded as the key GOBPs (Fig. S2, Table S1), and the AUCell scores of these key GOBPs were regarded as values of BP features (see Methods for details).

We took the UniCoord model trained by hECA data as a pre-trained model to analyze data that were not included in the training dataset. We used the model to represent cells in a hepatocellular carcinoma (HCC) dataset^38^. The dataset contains 56,721 cells from 46 distinct liver tumor samples (Fig. 4a-b). As shown in Fig. 4c-d, cells represented by the UniCoord model was less sensitive to the batch of samples. We then used UniCoord to annotate cell types of the HCC data (Fig. 4e). We found that the cell types unseen in hECA data were predicted as the related cell types (Fig. 4f). For example, cancer-associated fibroblasts (CAFs) were mainly annotated as smooth muscle cells as they share several important markers such as α-smooth muscle actin^39^. Besides, tumor-associated macrophages (TAMs) and tumor endothelial cells (TECs) were mainly annotated as their corresponding progenitor cell type, respectively. Interestingly, malignant cells were predicted as “others” which is a mixture of all cell types with small quantities in hECA. This suggested that the pre-trained UniCoord model successfully discovered that malignant cells were different from any cell type encoded in the model. Besides, B cells in the HCC data were also mainly predicted as “others”, suggesting that the state of these B cells was deviated from healthy B cells.

**Figure 4.**
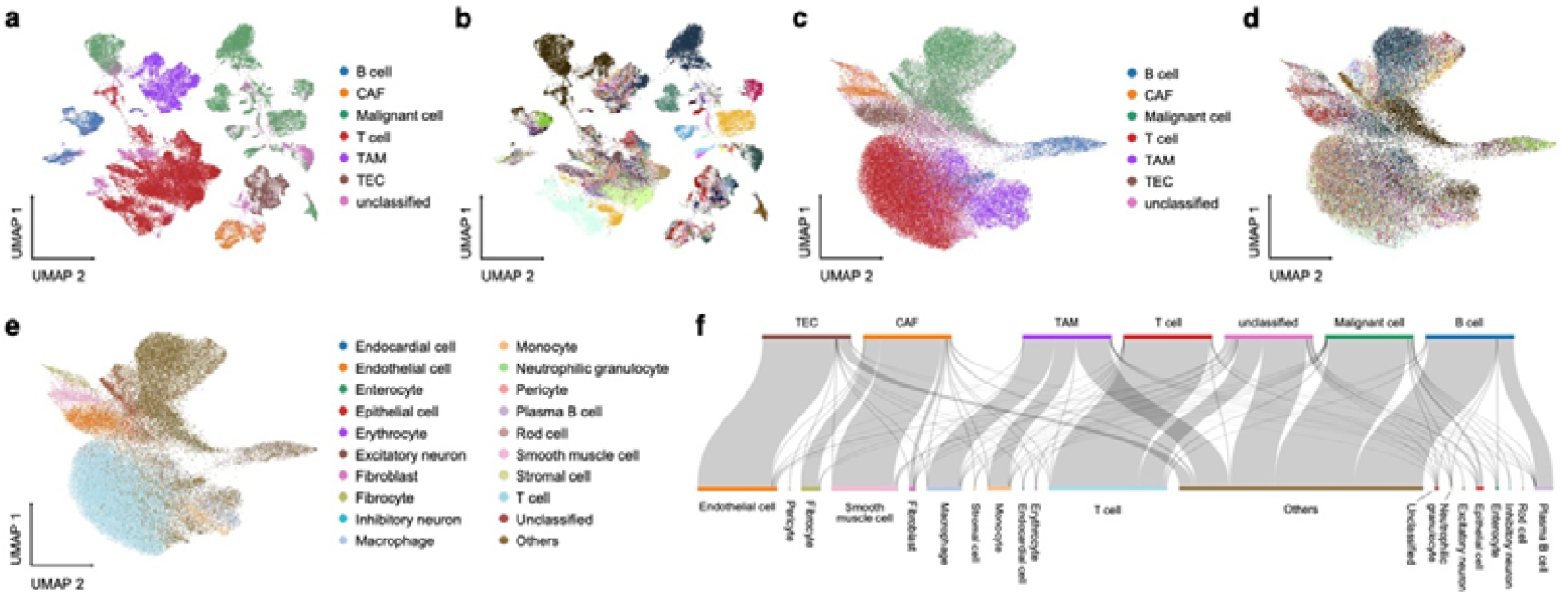
Analyze HCC data using the UniCoord model pre-trained by hECA data. (a-b) The UMAP plot showing the landscape of the HCC dataset, represented by PCA. Cells are colored by cell types (a) or sample ID (b). (c-e) The UMAP plot showing the landscape of the HCC dataset, represented by the pre-trained UniCoord model. Cells are colored by cell types (c), sample ID (d), and UniCoord-predicted cell types (e). (f) The relations between original labels (top) and cell types predicted by UniCoord (bottom). Protein coding genes in the dataset were adopted to perform the experiments. All genes were used for visualization all UMAP plots.

We investigated the BP features in the representation of the HCC data (Fig. S3, see Methods for details). We found that BP features such as B cell receptor signaling pathway, phagocytosis & recognition, and peptide cross-linking were highly activated in B cells. Connective tissue development, muscle cell development, and extracellular matrix organization were highly activated in CAFs. These BP features are highly related to the functions of the corresponding cell type. Malignant cells exhibited the lowest score of positive regulation of cell killing, which is consistent with their uncontrolled proliferation. The results showed feasibility of UniCoord as an interpretable pre-trained model for representing and decoding complex cell heterogeneities.

## Discussion

In this work, we presented UniCoord, a universal coordinate model that can represent cells with discrete and continuous features computationally derived from gene expression and/or metadata. We showed that UniCoord can efficiently capture the information in single-cell transcriptomic profiles. The resulting features can be used to well reconstruct the original transcriptional profiles and generate pseudo cells. The features can be either unsupervised or supervised by prior knowledge. The supervised features can be further interpreted for understanding cellular heterogeneities, and the unsupervised features can help extract information remaining to be characterized in current studies. Moreover, we demonstrated that UniCoord can serve as a pre-trained model that can be generalized to unseen data or cell types. These capabilities make UniCoord a powerful tool for analyzing the highly orchestrated functions and multifaceted heterogeneities of scRNA-seq data.

One major advantage of UniCoord is its capability to generate pseudo single-cell data by *in silico* reconfiguration, which can serve as a controlled generative model for data argumentation. This may be helpful to obtain transcriptomic profiles of desired cells or cell states that can hardly be observed experimentally. We demonstrated *in silico* reconfiguration by configuring sequencing platforms and cell types. We also showed that by *in silico* reconfiguration, UniCoord can reconstruct continuous trajectories from discrete data by interpolating the missing time points or unsampled spatial locations. This approach can help integrate datasets from different sources and generate pseudo cells to fill spatial, temporal or functional gaps in current data, both of which will contribute to the construction of cell atlases by unified organization and data argumentation. Furthermore, as an interpretable model, UniCoord is complemented with the fashionable large-scale pretrained models^40–42^. The results produced by these large models can be interpreted by UniCoord, and UniCoord can generate pseudo cells with high confidence to fit the huge demand of data for large model training.

It should be noted that the performance of UniCoord may be influenced by the quality and quantity of training data, particularly in smaller or less diverse datasets. Although UniCoord can interpolate data to fill gaps, the accuracy of the interpolated values could be limited in situations with high data sparsity or large gaps. UniCoord could benefit from the development of a more comprehensive and refined cell atlas that covers a wider range of cell types and aspects of cellular heterogeneities. Besides, the design of BP features can be further improved to enhance the clarity. In the future, UniCoord can be extended to more types of omics data to form a multi-scale framework for representing cellular complexity, and more applications such as measures of cellular functional distance and *in silico* perturbation of cell states can be further explored based on the framework.

## Methods

### The model of UniCoord

UniCoord was derived from the joint-VAE model^32^ with some refinements. The latent space of UniCoord differs from conventional VAE in two aspects: (a) the latent space of UniCoord is a combination of discrete and continuous dimensions, while conventional VAE only contain continuous dimensions; (b) each dimension in UniCoord could be physically interpretable if supervised by prior knowledge.

The gene expression level *x*_*i*_ of cell *i* could be modeled by a conditional distribution *p*(*x*_*i*_ | *ID*_*i*_, *UD*_*i*_, *IC*_*i*_, *UC*_*i*_), where *ID*_*i*_, *UD*_*i*_, *IC*_*i*_, and *UC*_*i*_ stand for interpretable discrete, unsupervised discrete, interpretable continuous and unsupervised continuous latent dimensions for cell *n*, respectively. *ID*_*i*_ and *IC*_*i*_ capture information corresponding to well defined features of the cell, such as the activity of a certain biological process, the differentiation stage of the cell, or clinical diagnoses of the cell’s donor. *UD*_*i*_ and *UC*_*i*_ capture complementary, yet unknown information in the data. *UD*_*i*_ and *UC*_*i*_ also play auxiliary roles that help the model reconstruct the original data. The mapping function *p* from these latent dimensions to expression levels is learned by training a neural network called decoder, and the posterior distribution of latent variables *q*(*ID*_*i*,_ *UD*_*i*_, *IC*_*i*_, *UC*_*i*_|*x*_*i*_) is learned by training another neural network called encoder.

### Model structure

The UniCoord model consists of an encoding module, a reparameterization module, and a decoding module. The encoding module processes the input data through three linear layers, with dimensions of 512, 256, and 128, respectively. The first and second linear layers have a dropout probability of 0.1. All three linear layers use the ReLU activation function. The output of the third linear layer was then transformed into two groups of parameters serving as the latent space. For continuous latent dimensions, two linear layers are used to map the output of third linear layer to mean and logarithm of variance, respectively. For each discrete latent dimension, a linear layer is used to map the output to its one-hot encoded value.

The reparameterization module samples from the latent space constructed by the encoder. The reparameterization trick can disentangle random variables with parameters and make back propagation algorithm possible. For continuous latent dimensions, we kept the conventional reparameterization trick used in VAE^31^. For discrete latent dimensions, we applied the Gumbel-softmax reparameterization^43^ for sampling.

The decoding module maps the samples provided by the reparameterization module back to the representation of the original data. The decoding model uses four linear layers to map the data from the dimensions of the latent samples to 128, 256, 512, and the dimensions of the original data, respectively. The second and third linear layers have a dropout probability of 0.1. All four linear layers use ReLU activation functions. The output of the decoding module is regarded as the generated data.

In all experiments of this study, the discrete dimensions were encoded by one-hot encoding. For all models with unsupervised latent dimensions, the number of unsupervised continuous dimensions was set as 50, the number of unsupervised discrete dimensions was set as 0.

### Loss functions

The loss function of UniCoord is composed of several parts, and each part gives the model a specific feature. In general,

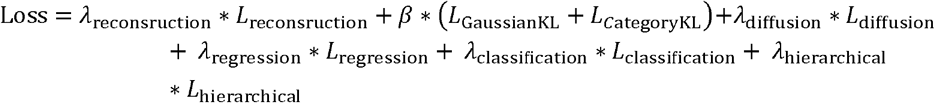

All *λ*s are hyper parameters that control the weight of each part, and details of each loss were introduced below.

#### Reconstruction loss

The reconstruction loss is the basic part of losses that make the model an auto-encoder. It is defined as the MSE between the reconstructed data ***x‰*** and original data ***x***:

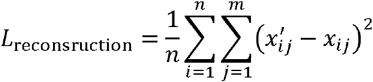

Where *n* is the number of cells, *m* is the number of genes, *x*_*lj*_ represents the expression level of gene *j* in cell *i*, and 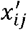 represents the reconstructed expression level of gene *j* in cell *i*.

#### KL divergence

KL divergence works as the regularization component that prevent over-fitting. For continuous dimensions, KL divergence constrains the posterior distribution to be closed to a standard normal distribution. For discrete dimensions, KL divergence constrains the posterior distribution to be closed to a uniform categorical distribution.

#### Diffusion loss

Some of the continuous dimensions can be defined as diffusion dimensions, playing the same roles as reductions in the diffusion map. We desired cells with similar scores in diffusion dimensions should also be similar in expression levels. So, we first constructed *k*-nearest neighbors for all cells and then calculated the average of the latent distribution of one cell’s neighbors. The average was inputted into the decoder to generate a reconstruction expression vector ***x*′′**. The diffusion loss is defined as the MSE between ***x*′′** and the original data ***x***.

#### Regression loss

The regression loss is the MSE loss between the original label and the mean parameter of the corresponding latent continuous dimension.

#### Classification loss

The classification loss is the cross entropy between the original label and the corresponding latent discrete dimension.

#### Hierarchical loss

The hierarchical loss is the cross entropy between two latent discrete dimensions that are designed to have hierarchical relationships. The descendant layer labels are first aggregate to ancestor labels following the designed relationship. Then the cross entropy between the aggregated labels and the model-generated ancestor labels is defined as the hierarchical loss.

### Model training

The “chunk_size” parameter is used to divide the training dataset into multiple parts for training in chunks. This is particularly useful for large datasets as it can reduce the memory and computation resources required for each iteration to improve training efficiency. By default, the “chunk_size” is set to 20000. The model is optimized using the Adam optimizer by default, and the default learning rate is 5e-4.

### Generation of BP features

The BP features are selected from GOBPs. To avoid outliers of enrichment analysis, we kept GOBP gene sets with gene number between 50 to 500, which resulted in 2,535 gene sets. Each of these gene sets was used to calculate an enrichment score for all cells in hECA data. Enrichment scores were calculated for each gene set across all cells in hECA data using the AUCell function in the SCENIC package^35^ with default parameters. Information entropy was then calculated for the AUCell scores of each gene set, and 10,000-fold permutation tests were performed to obtain *p*-values of the information entropy. We kept gene sets with *p*-values < 0.001 to train a random forest classifier to classify cell types in the hECA dataset annotated by the unified Hierarchal Annotation Framework (uHAF)^4^. The feature importance was evaluated using the feature_importance score of the classifier, and gene sets with the top 100 most important scores were selected to calculate the Pearson’s correlation coefficients between each other’s scores. Hierarchical clustering was performed to identify gene set groups with high correlations. We cut the hierarchical clustering to 30 groups and selected the one with the highest information entropy from each group. These 30 gene sets and their corresponding AUCell scores make up the BP features.

### scRNA-seq data analysis

Details of all scRNA-seq datasets used in this study can be found in Data Availability. The scRNA-seq data analysis was based on the python package Scanpy^44^. As UniCoord needed the normalized data, we applied sc.pp.normalize_total (target_sum = 1e4, exclude_highly_expressed = True), and sc.pp.log1p (default parameters) before feeding data into our model. For visualizing the landscape of single cells, we follow the standard analysis tutorial of Scanpy, and the data generated from UniCoord were also handled with the same procedure.

Analysis related to UniCoord was conducted with our python package, unicoord. Differential expressed genes and differential BP feature activities (Fig. S2) were detected using the tl.rank_genes_groups function in Scanpy.

## Data availability

We only used public datasets in this study. The hECA data can be downloaded through the Python package ECAUGT^45^. The mouse endothelial atlas dataset is available on the ArrayExpress database, with the Entrez accession number E-MTAB-8077. The mouse embryo trajectory data is available on the Gene Expression Omnibus (GEO) database with the accession number GSE122662. The human heart cell atlas data can be downloaded from its interactive website (http://xglab.tech/hahca). The HCC dataset is available on the GEO database with the accession number GSE151530.

## Code availability

The python package unicoord can be found in https://github.com/pluto-the-lost/unicoord.

## Acknowledgements

This work was partially supported by National Key R&D Program of China (grant 2021YFF1200901), National Natural Science Foundation of China (NSFC) (grants 62250005, 61721003 and 62103227), the CZI HCA Seed Network grant 2019-02444, and Tsinghua-Fuzhou Institute for Data Technology (TFIDT2021005). This publication is part of the Human Cell Atlas (www.humancellatlas.org/publications/).

## Author contributions

X.Z., L.W., H.G. and K.H. conceived the study. X.Z. and L.W. supervised the study. H.G. and K.H. design the model. H.G., K.H. and X.W. performed model training data analysis. S.C., Q.Y. and R.J. contributed to the improvement of the model and the interpretation of results. X.Z., L.W., H.G. and K.H. wrote the manuscript. All authors read and approved the final manuscript.

## Supplemental Materials

**Supplemental Figure 1.**
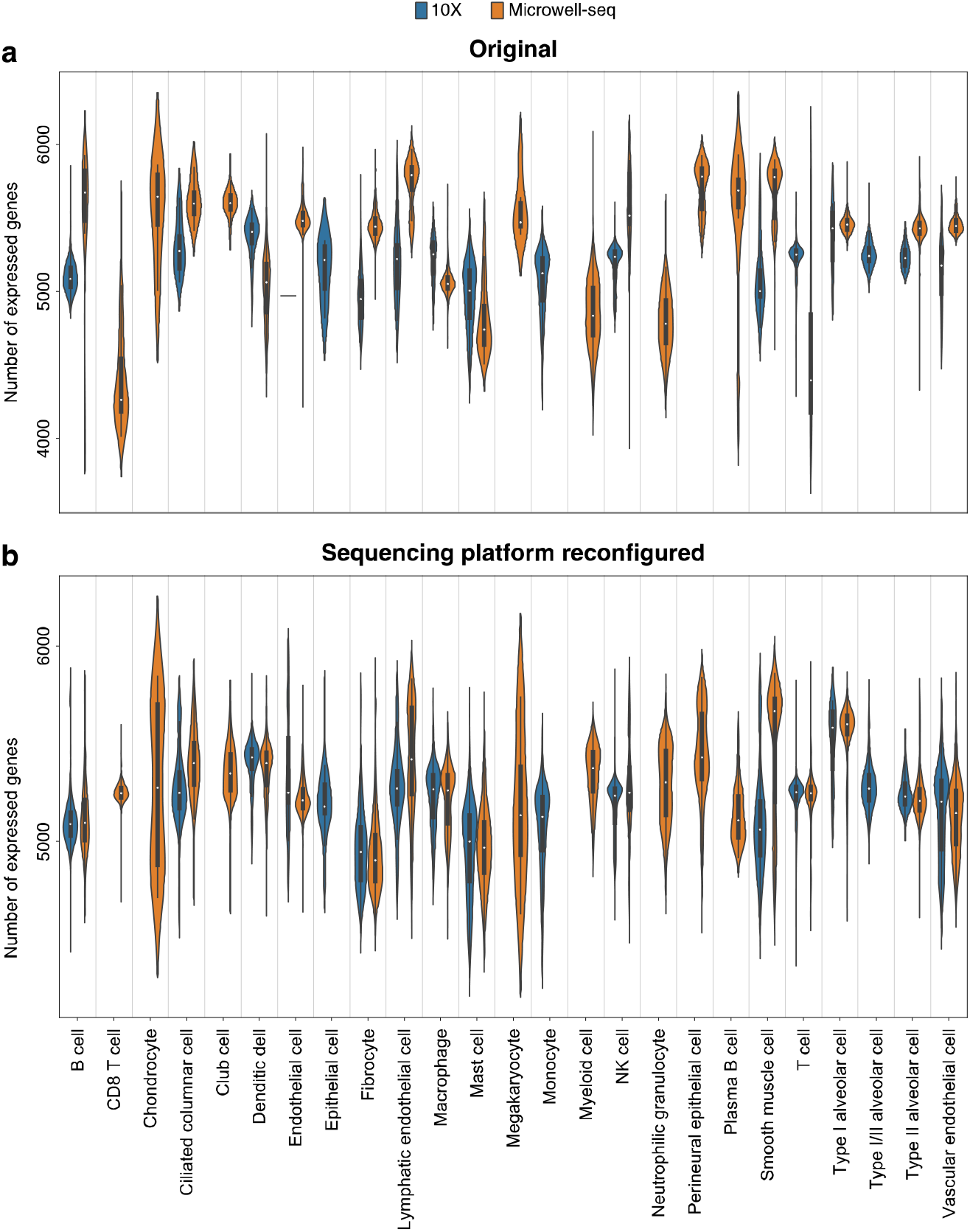
Distribution of the number of expressed genes with different sequencing platforms in the original cells (a) and cells with sequencing platformed reconfigured into 10X (b).

**Supplemental Figure 2.**
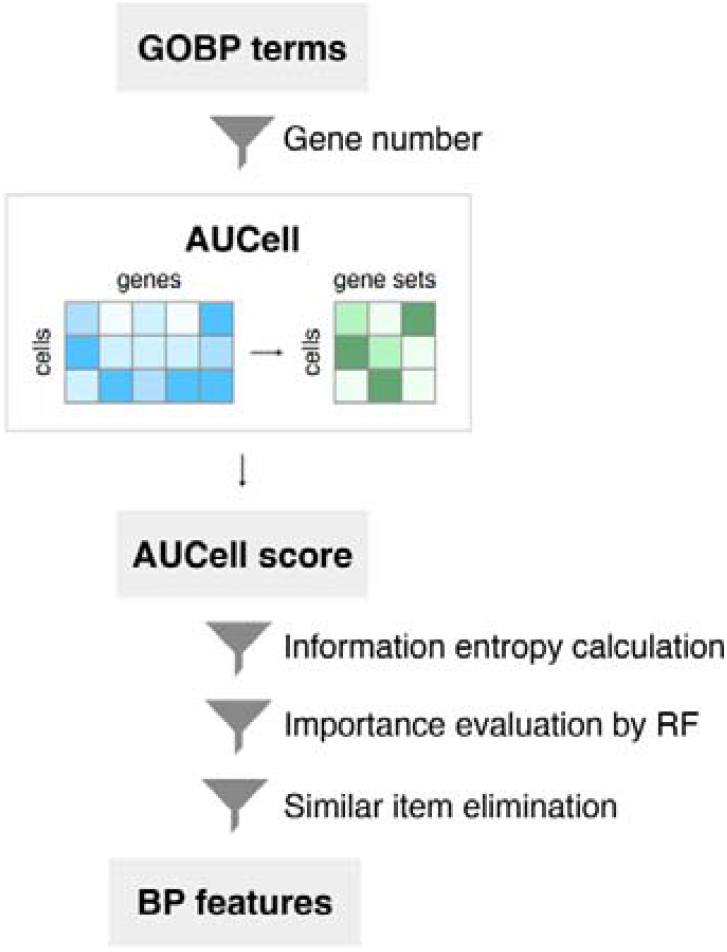
The generation of BP features.

**Supplemental Figure 3.**
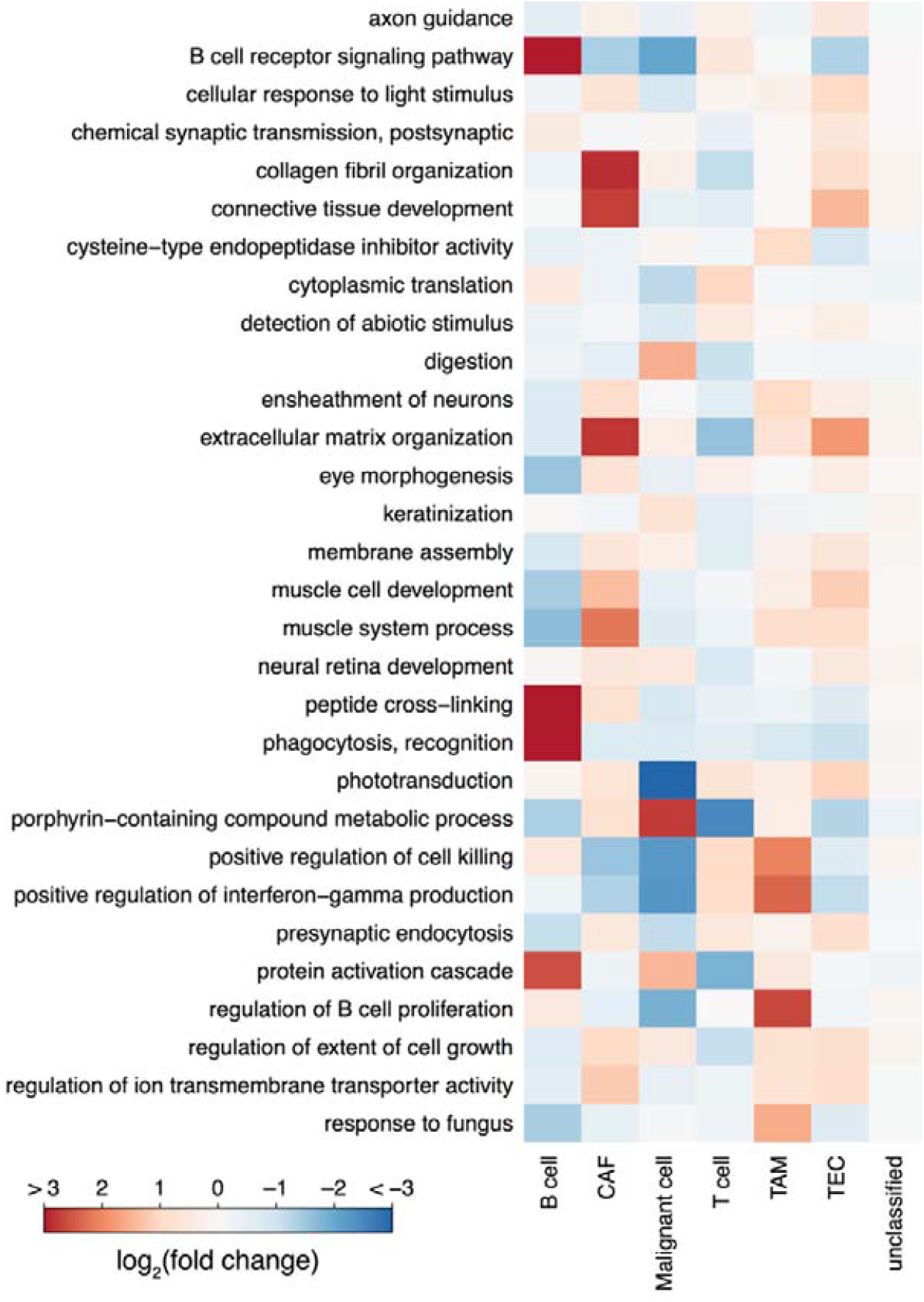
Fold changes of BP features of each cell type in the HCC dataset.

**Supplemental Table 1**. The list of BP features.

**Supplemental Movie 1**. UMAP plots of original and cells reconfigured with different zonation scores.

## References

1. Zeng, H. What is a cell type and how to define it? Cell 185, 2739–2755 (2022).

2. Regev, A. et al. The Human Cell Atlas. Elife 6, e27041 (2017).

3. HuBMAP Consortium et al. The human body at cellular resolution: the NIH Human Biomolecular Atlas Program. Nature 574, 187–192 (2019).

4. Chen, S. et al. hECA: The cell-centric assembly of a cell atlas. iScience 25, 104318 (2022).

5. Trapnell, C. et al. The dynamics and regulators of cell fate decisions are revealed by pseudotemporal ordering of single cells. Nat Biotechnol 32, 381–386 (2014).

6. Ji, Z. & Ji, H. TSCAN: Pseudo-time reconstruction and evaluation in single-cell RNA-seq analysis. Nucleic Acids Res 44, e117–e117 (2016).

7. Liu, Z. et al. Reconstructing cell cycle pseudo time-series via single-cell transcriptome data. Nat Commun 8, 22 (2017).

8. La Manno, G. et al. RNA velocity of single cells. Nature 560, 494–498 (2018).

9. Street, K. et al. Slingshot: cell lineage and pseudotime inference for single-cell transcriptomics. BMC Genomics 19, 477 (2018).

10. Saelens, W., Cannoodt, R., Todorov, H. & Saeys, Y. A comparison of single-cell trajectory inference methods. Nat Biotechnol 37, 547–554 (2019).

11. Satija, R., Farrell, J. A., Gennert, D., Schier, A. F. & Regev, A. Spatial reconstruction of single-cell gene expression data. Nat Biotechnol 33, 495–502 (2015).

12. Cang, Z. & Nie, Q. Inferring spatial and signaling relationships between cells from single cell transcriptomic data. Nat Commun 11, 2084 (2020).

13. Biancalani, T. et al. Deep learning and alignment of spatially resolved single-cell transcriptomes with Tangram. Nat Methods 18, 1352–1362 (2021).

14. Hao, M., Wei, L. & Zhang, X. Learning Spatially-Aware Representations of Transcriptomic Data via Transfer Learning. http://biorxiv.org/lookup/doi/10.1101/2022.09.23.509186 (2022) doi:10.1101/2022.09.23.509186.

15. Wei, R. et al. Spatial charting of single-cell transcriptomes in tissues. Nat Biotechnol 40, 1190–1199 (2022).

16. Diehl, A. D. et al. The Cell Ontology 2016: enhanced content, modularization, and ontology interoperability. J Biomed Semant 7, 44 (2016).

17. Osumi-Sutherland, D. et al. Cell type ontologies of the Human Cell Atlas. Nat Cell Biol 23, 1129–1135 (2021).

18. Sha, Y., Wang, S., Zhou, P. & Nie, Q. Inference and multiscale model of epithelial-to-mesenchymal transition via single-cell transcriptomic data. Nucleic Acids Research 48, 9505–9520 (2020).

19. Becker, W. R. et al. Single-cell analyses define a continuum of cell state and composition changes in the malignant transformation of polyps to colorectal cancer. Nat Genet 54, 985–995 (2022).

20. Hsiao, C. J. et al. Characterizing and inferring quantitative cell cycle phase in single-cell RNA-seq data analysis. Genome Res. 30, 611–621 (2020).

21. Li, C. et al. Single-cell transcriptomics–based MacSpectrum reveals macrophage activation signatures in diseases. JCI Insight 4, e126453 (2019).

22. Rood, J. E. et al. Toward a Common Coordinate Framework for the Human Body. Cell 179, 1455–1467 (2019).

23. Lopez, R., Regier, J., Cole, M. B., Jordan, M. I. & Yosef, N. Deep generative modeling for single-cell transcriptomics. Nature Methods 15, 1053–1058 (2018).

24. Stuart, T. et al. Comprehensive Integration of Single-Cell Data. Cell 177, 1888–1902.e21 (2019).

25. Chen, S. et al. Toward a unified information framework for cell atlas assembly. National Science Review 9, nwab179 (2022).

26. Bian, Z. et al. Deciphering human macrophage development at single-cell resolution. Nature 582, 571–576 (2020).

27. Zhang, M. et al. Spatially resolved cell atlas of the mouse primary motor cortex by MERFISH. Nature 598, 137–143 (2021).

28. Chen, L. et al. Multifaceted Spatial and Functional Zonation of Cardiac Cells in Adult Human Heart. Circulation 145, 315–318 (2022).

29. Conde, C. D. et al. Cross-tissue immune cell analysis reveals tissue-specific features in humans. Science 13 (2022).

30. Elmentaite, R., Domínguez Conde, C., Yang, L. & Teichmann, S. A. Single-cell atlases: shared and tissue-specific cell types across human organs. Nat Rev Genet (2022) doi:10.1038/s41576-022-00449-w.

31. Kingma, D. P. & Welling, M. Auto-Encoding Variational Bayes. (2013) doi:10.48550/ARXIV.1312.6114.

32. Dupont, E. Learning Disentangled Joint Continuous and Discrete Representations. in Advances in Neural Information Processing Systems (eds. Bengio, S. et al.) vol. 31 (Curran Associates, Inc., 2018).

33. Kalucka, J. et al. Single-Cell Transcriptome Atlas of Murine Endothelial Cells. Cell 180, 764–779.e20 (2020).

34. Schiebinger, G. et al. Optimal-Transport Analysis of Single-Cell Gene Expression Identifies Developmental Trajectories in Reprogramming. Cell 176, 928–943.e22 (2019).

35. Aibar, S. et al. SCENIC: single-cell regulatory network inference and clustering. Nat Methods 14, 1083–1086 (2017).

36. Ashburner, M. et al. Gene Ontology: tool for the unification of biology. Nat Genet 25, 25–29 (2000).

37. The Gene Ontology Consortium et al. The Gene Ontology resource: enriching a GOld mine. Nucleic Acids Research 49, D325–D334 (2021).

38. Ma, L. et al. Single-cell atlas of tumor cell evolution in response to therapy in hepatocellular carcinoma and intrahepatic cholangiocarcinoma. Journal of Hepatology 75, 1397–1408 (2021).

39. Han, C., Liu, T. & Yin, R. Biomarkers for cancer-associated fibroblasts. Biomark Res 8, 64 (2020).

40. Theodoris, C. V. et al. Transfer learning enables predictions in network biology. Nature (2023) doi:10.1038/s41586-023-06139-9.

41. Cui, H. et al. scGPT: Towards Building a Foundation Model for Single-Cell Multi-omics Using Generative AI. http://biorxiv.org/lookup/doi/10.1101/2023.04.30.538439 (2023) xdoi:10.1101/2023.04.30.538439.

42. Hao, M. et al. Large Scale Foundation Model on Single-cell Transcriptomics. http://biorxiv.org/lookup/doi/10.1101/2023.05.29.542705 (2023) xdoi:10.1101/2023.05.29.542705.

43. Jang, E., Gu, S. & Poole, B. Categorical Reparameterization with Gumbel-Softmax. (2016) doi:10.48550/ARXIV.1611.01144.

44. Wolf, F. A., Angerer, P. & Theis, F. J. SCANPY: large-scale single-cell gene expression data analysis. Genome Biol 19, 15 (2018).

45. Chen, Y. et al. Protocol for profiling cell-centric assembled single-cell human transcriptome data in hECA. STAR Protocols 3, 101589 (2022).

